# Chromosome Ageing Occurs at the Primordial Follicle Stage in both Mouse and Human Oocytes

**DOI:** 10.1101/2025.07.17.663489

**Authors:** Matilda Bui, Apiwat Moolnangdeaw, Hannes Becher, Shelagh Boyle, Vlastimil Srsen, Yvonne L. Odey, Richard A. Anderson, Evelyn E. Telfer, Ian R. Adams

## Abstract

Inherited aneuploidy is one of the most common causes of human genetic disease affecting approximately 20% of all human embryos and is the major cause of non-conception in older women. Inherited aneuploidies typically arise from meiotic chromosome segregation errors and are strongly affected by maternal age. Cohesin, a protein complex that mediates cohesion between sister chromatids, becomes depleted from oocyte chromosomes with age in mice, potentially contributing to the high rate of aneuploidy in older mouse oocytes. However, it remains unclear at which stage of oogenesis ageing affects chromatid cohesion in mice and whether ageing similarly affects cohesin in human oocytes. Here we use fluorescence in situ hybridisation to assess at which stages of oogenesis ageing alters chromatid separation in mice and in humans. We show that chromatid separation increases with age while oocytes remain dormant in primordial follicles in both mice and humans and that this effect of age on chromatid separation is detectable in fertile women of child-bearing age. Furthermore, we show that this age-dependent increase in chromatid separation is accompanied by age-dependent depletion of the acetylated SMC3-marked cohesive subpopulation of cohesin from oocytes at the dormant primordial follicle stage in both mice and humans. These data suggest that ageing impacts on oocyte chromosomes while they remain dormant in primordial follicles, and that this aspect of oocyte chromosome ageing is shared in both mice and humans. Improving our understanding of these pathways may allow strategies to slow or prevent this aspect of oocyte ageing to be developed.

## Introduction

Approximately 20% of human embryos, corresponding to around 30-40 million conceptions annually, inherit the wrong number of chromosomes^1–6^. These inherited constitutive aneuploidies, a leading cause of conception failure, miscarriage and developmental disorders such as Down syndrome, arise primarily through meiotic chromosome segregation errors during parental gametogenesis. These errors occur far more frequently during oogenesis than spermatogenesis, and are strongly influenced by maternal age with most errors in older human oocytes arising during the first meiotic division^1–6^. Several mechanisms have been proposed to explain this maternal age effect, including a ‘production line’ hypothesis that higher quality oocytes ovulate earlier in adult life^7^, ovulation-induced damage^8^, and age-dependent deterioration of meiotic chromosomes^9–14^, the chromosome segregation machinery^15–17^ or checkpoints^18–20^. However, the mechanistic causes of age-dependent meiotic chromosome segregation errors during human oogenesis remain to be fully established.

In mammals, oocytes typically arrest in meiosis at the dictyate stage and, before or around the time of birth, interact with surrounding somatic cells to form primordial follicles^1–4,21^. These follicles establish the finite pool of oocytes that must sustain the female reproductive lifespan. In humans, primordial follicles can remain dormant for decades before being activated to grow^1–4,21^, and thus the length of time that an oocyte has remained dormant at the primordial follicle stage increases with age (**Figure 1A**)^1–4,21^. Upon primordial follicle activation, the oocyte grows and matures as the follicle progresses through primary and secondary stages. Around ovulation, fully-grown mature oocytes within antral follicles exit dictyate and subsequently progress through two rounds of meiotic chromosome segregation^1–4,21^. Much of our knowledge about the effects of age on human oocytes comes from fully-grown oocytes obtained either after ovulation during assisted reproduction procedures, or after growth and maturation in vitro^11,17,22–24^. Understanding which stages of postnatal oogenesis are affected by age both in the non-growing primordial follicle and at later stages, and how ageing impacts on oocyte meiotic chromosome segregation, is key to developing strategies to reduce maternally-transmitted aneuploidies.

**Figure 1.**
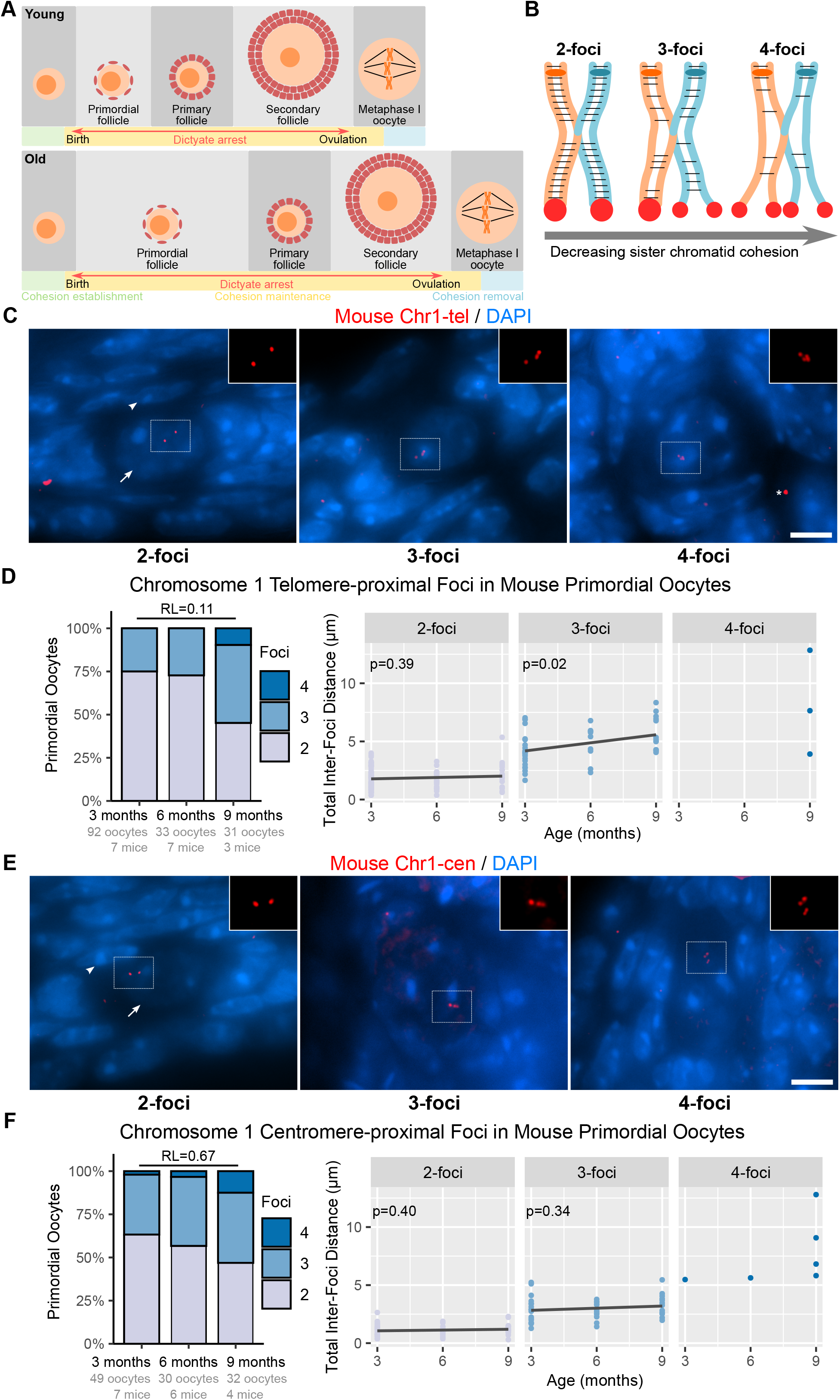
Fluorescence in situ hybridisation detects age-dependent changes in chromatid separation in mouse dictyate stage oocytes in dormant primordial follicles. **A**. Schematic diagram showing how age affects the duration of stages of oogenesis in mice. Morphology of oocytes (dark orange, nucleus/chromosomes; light orange, cytoplasm) and granulosa cells (red) are shown, regulation of cohesion is indicated by colours in the timeline. **B**. Schematic diagram showing how reduced sister chromatid cohesion can affect chromatid separation within bivalent mouse chromosomes. Orange/blue, homologous chromosomes; ellipses, centromeres; black bars, cohesion; red, telomere-proximal FISH signals. More complex configurations are possible, chiasma may also influence chromatid separation in nearby regions. **C, E**. FISH for chromosome 1 telomere-proximal (Chr1-tel, C) and centromere-proximal (Chr1-cen, E) fosmids (red) in primordial follicles in mouse ovary sections. FISH signals are 3D maximum projections, DNA is stained with DAPI (blue). Arrow, oocyte; arrowhead, flattened granulosa cell; asterisk, non-specific signal; dashed rectangle, insets (FISH channel only). Scale bar 5 µm. **D, F**. Quantification of Chr1-tel (D) and Chr1cen (F) FISH foci in mouse primordial oocytes. Left panels, distribution of oocytes with the indicated number of foci. RL, relative likelihood that a logit model not incorporating age fits the foci number distribution better than one that does. Right panel, total Chr1-tel (D) and Chr1-cen (F) inter-foci distance within 2-foci, 3-foci and 4-foci mouse primordial oocytes groups against age. Black lines, locally weighted scatterplot smoothing trends; p-values, linear mixed-effect models.

Studies in mice suggest that age-dependent loss of chromosome-associated cohesin contributes to aneuploidy in older oocytes^9,10,17,25–27^. Cohesin is a multi-subunit protein complex with multiple roles in chromosome biology, including mediating cohesion between sister chromatids^28–30^. During meiosis, sister chromatid cohesion maintains the chiasma, the physical links between homologous chromosomes that result from crossover recombination, between chromosome arms and also the physical links between sister chromatids at their centromeres^17,25,26,31^. Step-wise removal of cohesin first from chromosome arms and then from centromeres allows ordered segregation of homologous chromosomes then sister chromatids during the first and second rounds of meiotic chromosome segregation respectively^26,31–34^. Cohesins are loaded onto meiotic chromosomes during fetal stages of oocyte development then maintained throughout the oocytes’ prolonged postnatal dictyate arrest^26,35,36^. The amount of the REC8-cohesin subunit associated with meiotic chromosomes in fully-grown metaphase I mouse oocytes, though not the total amount of REC8 in these cells, dramatically decreases with increasing age^9,10^. Age-dependent depletion of chromosome-associated REC8 in older growing oocytes is also evident at the earlier preantral follicle stage of oogenesis^9^. This age-dependent deterioration of chromosome-associated cohesin likely contributes to the loss and terminalisation of chiasma, increased sister kinetochore separation and increased chromosome missegregation evident in fully-grown metaphase I oocytes from older mice^7,9,10^. However, it is not clear how ageing causes deterioration of chromosome-associated cohesin in mouse oocytes, or even at which stage of oogenesis age-dependent depletion of cohesin occurs.

In contrast to mice, direct evidence that chromosome-associated cohesin declines with age in human oocytes remains limited. Meiotic chromosome mis-segregation patterns^5^, premature resolution of bivalents during the first meiotic division^23^, and increased sister kinetochore separation in the first and second meiotic division^11,17,22,37,38^ are all consistent with reduced levels of cohesion in older human oocytes. Furthermore, immunostaining for the SMC1β cohesin subunit in dictyate oocytes within ovary tissue sections has been reported to be lower in older women^39^. However, it is less clear whether anti-cohesin immunostaining of ovary tissue sections reflects changes in the chromosome-associated population of cohesin or reduced cohesin function and, in contrast to mice, age-dependent depletion of REC8-cohesin from isolated oocyte chromosomes has not been detected at metaphase I^24^. Therefore, at present it is not clear whether chromosome-associated cohesins are functionally affected by ageing in human oocytes similarly to mice.

We recently demonstrated^40^ that postnatal mouse oocytes possess a regulatory pathway that specifically affects the subpopulation of chromosome-associated cohesin that mediates sister chromatid cohesion and marked by acetylation of the SMC3 subunit (AcSMC3)^41–44^. This study^40^ suggests only a minor fraction of chromosome-associated REC8 is involved in cohesion. Therefore, we sought to investigate whether analysis of sister chromatid cohesion and the cohesive subpopulation of cohesin might reveal commonalities in mice and humans in when and how ageing affects meiotic chromosomes in postnatal oocytes.

## Results

### Mouse oocytes exhibit chromosome ageing at the primordial follicle stage

Previous studies suggest that metaphase I oocyte chromosomes from older mice contain less REC8-cohesin, potentially contributing to age-dependent oocyte aneuploidy in this species^7,9,10,23^. To investigate at which stage of mouse oogenesis age affects chromatid cohesion, we developed a fluorescence in situ hybridisation (FISH) approach to assess chromatid separation at earlier stages of oogenesis. Studies using FISH in somatic cells and in *Drosophila* oocytes have shown that decreased levels of cohesin and cohesion results in increased numbers of discrete FISH foci detected at a locus (**Figure 1B**) and/or distances between these foci^44–47^. Therefore, we reasoned that FISH may similarly allow age-dependent depletion of cohesion in dictyate mammalian oocytes to be detected.

Chromatid cohesion is differentially regulated on chromosome arms and centromeres in meiosis I, with centromeric cohesion being more protected^48–50^. Therefore, we designed fosmid probes to label regions proximal to each of the centromere (Chr1-cen) and telomere (Chr1-tel) of mouse chromosome 1 (**Supplementary Figure S1**). Both probes were verified as detecting the desired chromosomal regions using FISH with chromosome 1 paints on mitotic chromosome spreads (**Supplementary Figure S1**). FISH on sections of fixed, wax-embedded adult mouse ovary sections using fosmid probes directly labelled with fluorophores preserved tissue morphology sufficiently well to allow dictyate oocytes and follicular stages to be identified, and discrete nuclear FISH signals to be detected over background (**Figure 1C**). Some non-specific signals were also sometimes visible, particularly in cytoplasmic or inter-cellular regions (**Figure 1C**). Dictyate stage meiotic oocytes contain four copies of each chromosomal locus and, as expected, typically contained 2, 3 or 4 discrete nuclear foci for these fosmids (**Figure 1C**). Thus, FISH on ovary sections is a viable method to directly assess chromatid organisation in postnatal mammalian oocytes.

We next investigated whether the age-dependent depletion of REC8-cohesin and chromatid cohesion that has occurred in fully-grown metaphase I mouse oocytes^7,9,10,18^ might reflect an age-dependent loss occurring earlier in oogenesis while dictyate oocytes remain dormant in primordial follicles. As REC8 becomes undetectable in mouse metaphase I oocyte chromosome spreads at around 6 months of age^9,10^, we used mice between 3 and 9 months old to span this age range. 73% of oocytes within primordial follicles (hereafter termed primordial oocytes) in ovary sections from 3month-old mice have 2 visible discrete Chr1-tel foci, with the remaining primordial oocytes having 3 foci (**Figure 1D**). However, the proportion of primordial oocytes with 2 Chr1-tel foci decreases to 45% by 9 months of age, and 10% of primordial oocytes now had 4 Chr1-tel foci (**Figure 1D**). Incorporating age as an effect in a logit model is 9 times more likely to explain the number of Chr1-tel foci in mouse primordial oocytes than a null model incorporating random differences between individual mice (**Figure 1D**). Therefore, increasing age is associated with increasing chromatid separation at the telomere-proximal region of chromosome 1 within mouse primordial oocytes.

To determine whether increasing age correlates with greater inter-chromatid distances in primordial oocytes, we grouped primordial oocytes by the number of Chr1-tel foci detected, then measured the total inter-foci distance for each oocyte. No strong age-dependent change was detected in total inter-foci distance in primordial oocytes with 2 visible discrete Chr1-tel foci (**Figure 1D**), however the total inter-foci distance for primordial oocytes containing 3 Chr1-tel foci, increases from a median of 4.0 µm at 3-months-old to a median of 5.1 µm at 9-months-old. (**Figure 1D**). This difference between 2-foci and 3-foci primordial oocytes effect may reflect distinct age-related changes in inter-homolog and inter-sister chromatid distances (**Figure 1B**).

FISH does not distinguish between chromatids from homologous chromosomes and sister chromatids. However, pairwise distances between Chr1-tel foci fitted bimodal distributions consistent with mouse primordial oocytes having two distinct populations of pairwise inter-chromatid distances (**Supplementary Figure S2**). These two populations potentially represent closer inter-sister and more separate inter-homolog distances. The inter-chromatid distances for both these populations increased with age in mouse primordial oocytes (**Supplementary Figure S2**). As the distances between foci in 2-foci primordial oocytes did not show a strong age-dependent effect (**Figure 1D**), these FISH data are consistent with first inter-sister chromatid distances, then inter-homolog distances, increasing with age at this chromosomal region.

We next investigated how age might affect chromatid separation at the centromere-proximal Chr1-cen region in mouse primordial oocytes (**Figure 1E**). 63% of primordial oocytes from 3-month-old mice have 2 visible discrete Chr1-cen foci and only 2% have 4 Chr1-cen foci (**Figure 1F**). Although the proportion of primordial oocytes with 4 Chr1-cen foci increases to 12% in 9-month-old mice, incorporating age does not strongly improve a logit model for Chr1-cen foci number (**Figure 1F**). Neither could we detect a strong age-dependent change in total Chr1-cen inter-foci distances in mouse primordial oocytes (**Figure 1F**). Therefore, although increasing sample size or age range may allow age-dependent changes at the Chr1-cen region to be detected in mouse primordial oocytes, chromatid separation at this centromere-proximal region of chromosome 1 seems to be less sensitive to age than the telomere-proximal region.

We next assessed whether any effects of age on chromatid separation in primordial oocytes persist as folliculogenesis progresses to the primary follicle stage. The number of Chr1-tel foci in oocytes within primary follicles (hereafter termed primary oocytes) is strongly affected by age with 75% of 3-month-old primary oocytes, but only 34% of 9-month-old primary oocytes, having 2 visible discrete Chr1-tel foci (**Supplementary Figure S3**). Primary oocytes with 4 Chr1-tel foci concomitantly increases from 3% to 17% between these ages, and incorporating age makes a logit model 3000 times more likely to fit the Chr1-tel foci number data for primary oocytes (**Supplementary Figure S3**). Furthermore, total inter-chromatid distance amongst 3-foci primary oocytes, but not 2-foci primary oocytes, increases with increasing age (**Supplementary Figure S3**). Chr1-cen was not strongly affected by age in primordial oocytes, and we could not detect age-dependent effects on chromatid separation at this region in primary oocytes either (**Supplementary Figure S3**). Thus, the age-dependent changes in chromatid separation that we detected in mouse primordial oocytes appear to persist and be propagated to the next stage of folliculogenesis.

### Human oocytes also exhibit chromosome ageing at the primordial follicle stage

We next tested whether human oocytes might similarly experience chromosome ageing while they remain dormant within primordial follicles. Given the impact of trisomy 21 aneuploidies^1–5^, and the high rate of meiosis I chromosome segregation errors for this chromosome with advanced maternal age^5^, we used a fosmid probe directed against a telomere-proximal region of chromosome 21 (Chr21-tel) (**Supplementary Figure S1**). Sections of ovarian cortical tissue containing primordial and early growing follicles were obtained from two groups of patients: young childhood cancer patients (< 18 years old) undergoing fertility preservation prior to cancer treatment, and adult women (≥ 18 years old) undergoing elective Caesarean section. FISH using the Chr21-tel probe on human ovary tissue (**Figure 2A**) shows that 69% of human primordial oocytes have 2 visible discrete Chr1-tel foci, 27% have 3 and and 4% have 4 Chr1-tel foci in the young patient group. However, adult women exhibit significantly more Chr1-tel foci in their primordial follicles, with only 22% having 2 Chr1-tel foci and 40% having 4 Chr21-tel foci (**Figure 2B**). Incorporating age group as an effect in a logit model is 400 times more likely to explain the number of Chr21-tel foci in human primordial oocytes (**Figure 2B**). Even within the adult women group, primordial oocytes in older (33-40 years old) adults have more Chr21-tel FISH foci than younger (25-32 years old) adults, and this age-dependent effect on adult oocyte chromosomes persists into the primary oocyte stage (**Supplementary Figure S4**). Therefore, at least some of the effect of age on chromatid separation occurs in primordial oocytes of adult women during the child-bearing age range. Total inter-chromatid distances for Chr21-tel also significantly increases with increasing age amongst human primordial oocytes with 3 discrete foci, but not amongst those with 2 discrete foci (**Figure 2B**). In contrast, Chr21-tel inter-foci distances in primordial follicle granulosa cells are not significantly affected by age (**Supplementary Figure S5**) suggesting that the age-dependent effects detected in primordial oocytes are unlikely to represent an artefact introduced during tissue processing. Thus, similarly to its effects on a telomere-proximal region of chromosome 1 in mice, ageing also affects chromatid organisation at a telomere-proximal region of chromosome 21 in human oocytes while they remain dormant at the primordial follicle stage.

**Figure 2.**
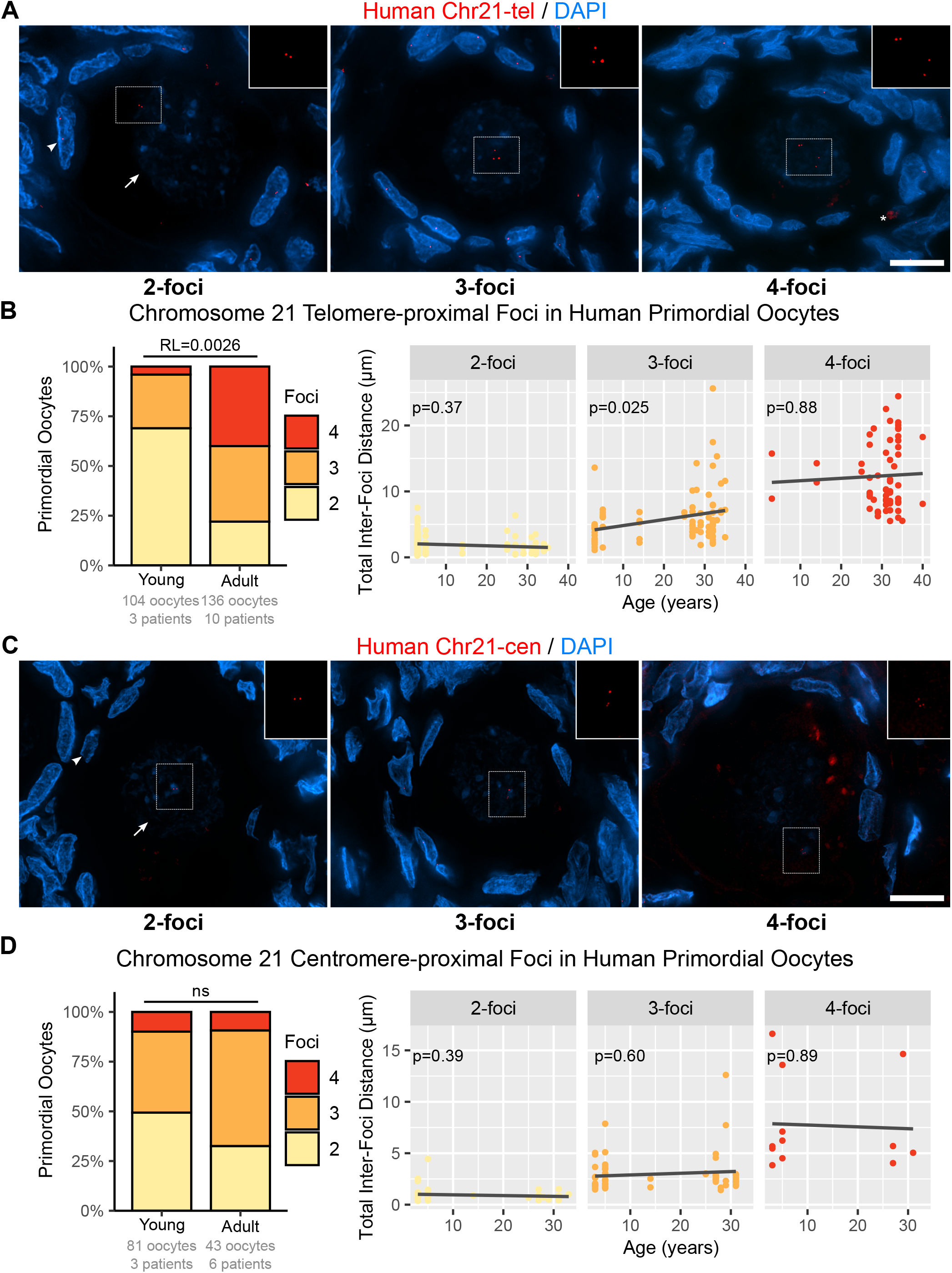
Fluorescence in situ hybridisation detects age-dependent changes in chromatid separation in human oocytes in primordial follicles. **A, C**. FISH on human ovary sections using Chr21 telomere-proximal (Chr21-tel, A) and Chr21 centromere-proximal (Chr21-cen, C) fosmid probes (red). DNA is counterstained with DAPI (blue), images are 3D maximum projections. Arrow, oocyte; arrowhead, flattened granulosa cell; asterisk, non-specific signal; dashed rectangle, insets (FISH channel only). Scale bar 5 µm. **B, D**. Quantification of Chr21-tel (B) and Chr21-cen (D) FISH foci in human primordial oocytes. Left panel, distribution of oocytes with the indicated number of foci. Young patients, 3-14 years old; adult women, 25-40 years old. RL, relative likelihood that a logit model not incorporating age fits the foci number distribution better than one that does. Right panel, total Chr21-tel (B) and Chr21-cen (D) inter-foci distance within 2-foci, 3-foci and 4-foci human primordial oocytes groups against age. Black lines, locally weighted scatterplot smoothing trends; p-values, linear mixed-effect models.

Similarly to the mouse FISH data, pairwise distances between Chr21-tel foci fitted bimodal distributions consistent with human primordial oocytes having two distinct populations of pairwise inter-chromatid distances, potentially representing inter-sister and inter-homolog distances (**Supplementary Figure S2**). The inter-chromatid distances for both these populations, but not distances between foci in 2-foci primordial oocytes, increased with age in humans (**Supplementary Figure S2**). Therefore, like Chr1-tel in mice, the ageing may affect inter-sister chromatid distances first, then inter-homolog distances, at the telomere-proximal region of chromosome 21 in humans.

We next investigated whether our findings from the telomere-proximal region of chromosome 21 might extend to other chromosomal regions in the human genome. Trisomies for chromosome 13 and 16 are implicated in miscarriage and Patau syndrome respectively^1–5^, and like chromosome 21, both these chromosomes exhibit meiosis I chromosome segregation errors at advanced maternal age^5^. The telomere-proximal region of chromosome 13 (Chr13-tel, **Supplementary Figure S1**) shows a significant increase in the proportion of primordial oocytes containing 4 visible discrete Chr13-tel foci, from 8% in young ovaries to 37% in adult ovaries, with the proportion of primordial oocytes containing 2 Chr13-tel foci decreasing from 56% to 25% across the same age range (**Figure 3**). However, total inter-foci distances at Chr13-tel does not significantly increase with age in human primordial oocytes (**Figure 3**). In contrast, while the number of visible discrete foci for the telomere-proximal region of chromosome 16 (Chr16-tel, **Supplementary Figure S1**) does not significantly change between young patient and adult women groups (**Figure 3**), total inter-foci distance at this chromosomal region for each of the 2-foci, 3-foci and 4-foci primordial oocyte categories all significantly increase with age (**Figure 3**). The strong age-dependent increase in total inter-foci distance for Chr16-tel is specific to oocytes as no age-dependent changes in inter-foci distances within the granulosa cells surrounding primordial follicles could be detected in these ovary sections (**Supplementary Figure S5**). Therefore, although telomere-proximal regions of chromosomes 13, 16 and 21 all exhibit age-dependent increases in chromatid separation in human primordial oocytes, there may be some differences in chromosome organisation or sister chromatid cohesion, or how these are affected by age, between these different chromosomal regions.

**Figure 3.**
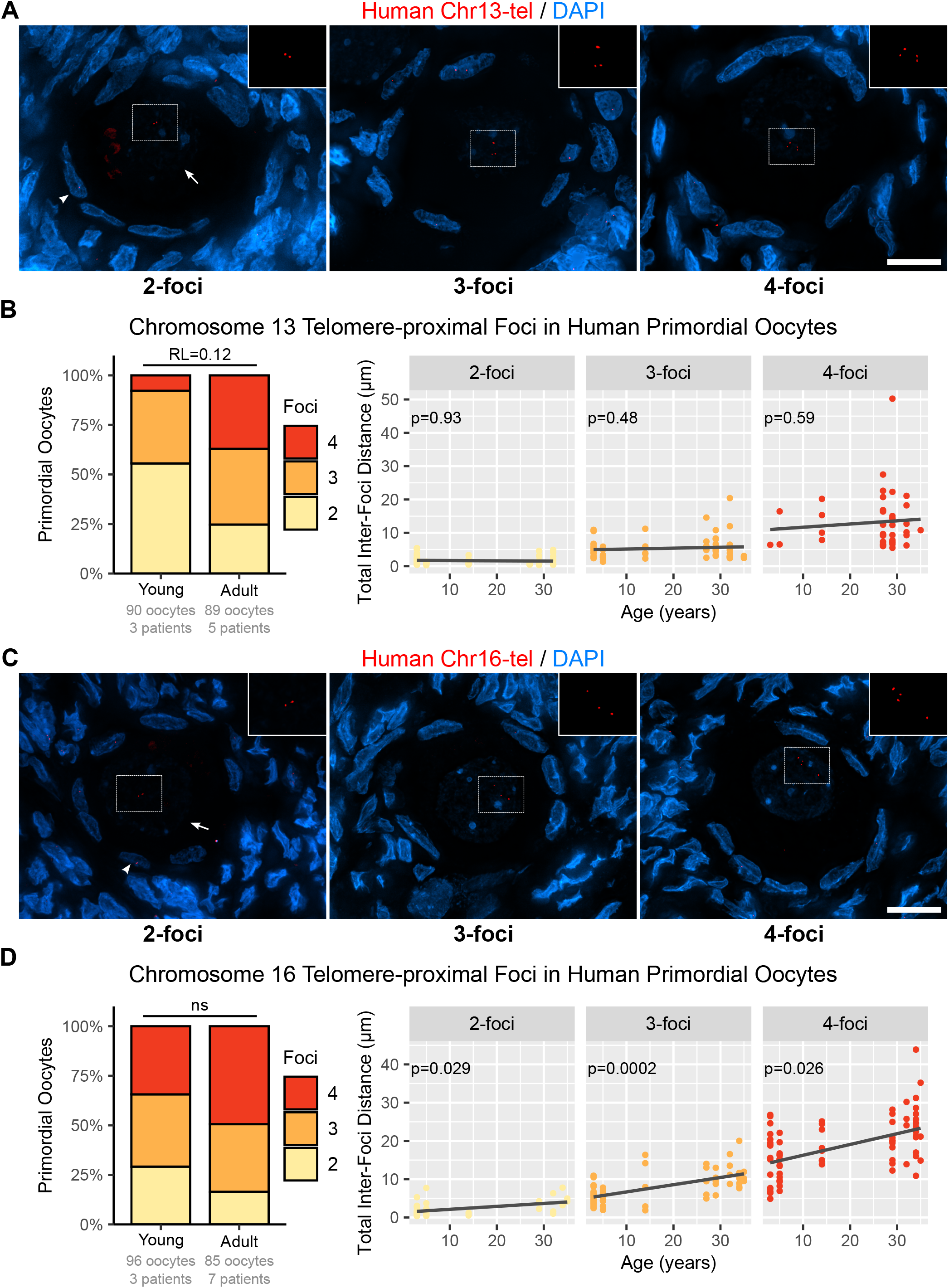
Chromatid separation at telomere-proximal regions of chromosomes 13 and 16 increase with age in human oocytes in primordial follicles. **A, C**. FISH on human ovary sections using Chr13 telomere-proximal probe (Chr13-tel, A) or a Chr16 telomere-proximal (Chr16-tel, B) fosmid probes (red). DNA is counterstained with DAPI (blue), images are 3D maximum projections. Arrow, oocyte; arrowhead, flattened granulosa cell; dashed rectangle, insets (FISH channel only). Scale bar 5 µm. **B, D**. Quantification of Chr13-tel (C) and Chr16-tel (D) FISH foci in human primordial oocytes. Left panel, distribution of oocytes with the indicated number of foci. Young patients, 3-14 years old; adult women, 25-36 years old. RL, relative likelihood that a logit model not incorporating age fits the foci number distribution better than one that does. Right panel, total Chr13-tel (C) and Chr16-tel (D) inter-foci distance within 2-foci, 3-foci and 4-foci human primordial oocytes groups against age. Black lines, locally weighted scatterplot smoothing trends; p-values, linear mixed-effect models.

We also assessed whether, like mouse, centromere-proximal regions of a human chromosome might be less sensitive to chromosome ageing in primordial oocytes. A FISH probe directed against human chromosome 21 (Chr21-cen, **Supplementary Figure S1**) does not exhibit a significant detectable change in the number of visible discrete Chr21-cen foci in primordial oocytes between young and adult ovaries (**Figure 2D**). Neither does Chr21-cen total inter-foci distance increase with age in human primordial oocytes (**Figure 2D**). Therefore, although increasing sample size or age range may allow an age-dependent change in chromatid separation at the Chr21-cen to be detected in human primordial oocytes, age-dependent effects on chromatid separation seem to be more restricted at this centromere-proximal region than at the three telomere-proximal regions tested.

### Acetylated SMC3-containing cohesin is depleted with age from mouse and human primordial oocytes

We next investigated whether these age-dependent increases in chromatid separation in mouse and human primordial oocytes might reflect age-dependent depletion of cohesin occurring in these cells. We previously reported that the AcSMC3-containing subpopulation of cohesin may better reflect the amount of sister chromatid cohesion in postnatal mouse oocytes than the REC8-containing population^40^. However, it is not known whether chromosome-associated AcSMC3 in oocytes is affected by ageing. Therefore, we first tested whether AcSMC3, like REC8, is depleted in metaphase I oocyte chromosomes from older mice. As reported previously^40^, AcSMC3 is enriched on the chromosome axes in chromosome spreads from young metaphase I mouse oocytes (**Figure 4**). Furthermore, similarly to REC8^9,10^, AcSMC3 immunostaining is significantly depleted from older metaphase I oocyte chromosomes, with oocytes from 12-month old mice containing 2.2-fold less chromosome-associated AcSMC3 than those from 3-month old mice (**Figure 4**). Anti-AcSMC3 immunostaining remains detectable in metaphase I oocyte chromosome spreads from 12-month old mice, consistent with oocytes having sufficient sister chromatid cohesion to maintain the structure of the bivalent (**Figure 4**) and prevent high levels of aneuploidy at this age^9,10,23^. Thus, the AcSMC3-containing subpopulation of cohesin implicated in chromatid cohesion becomes depleted from metaphase I oocyte chromosomes as mice age.

**Figure 4.**
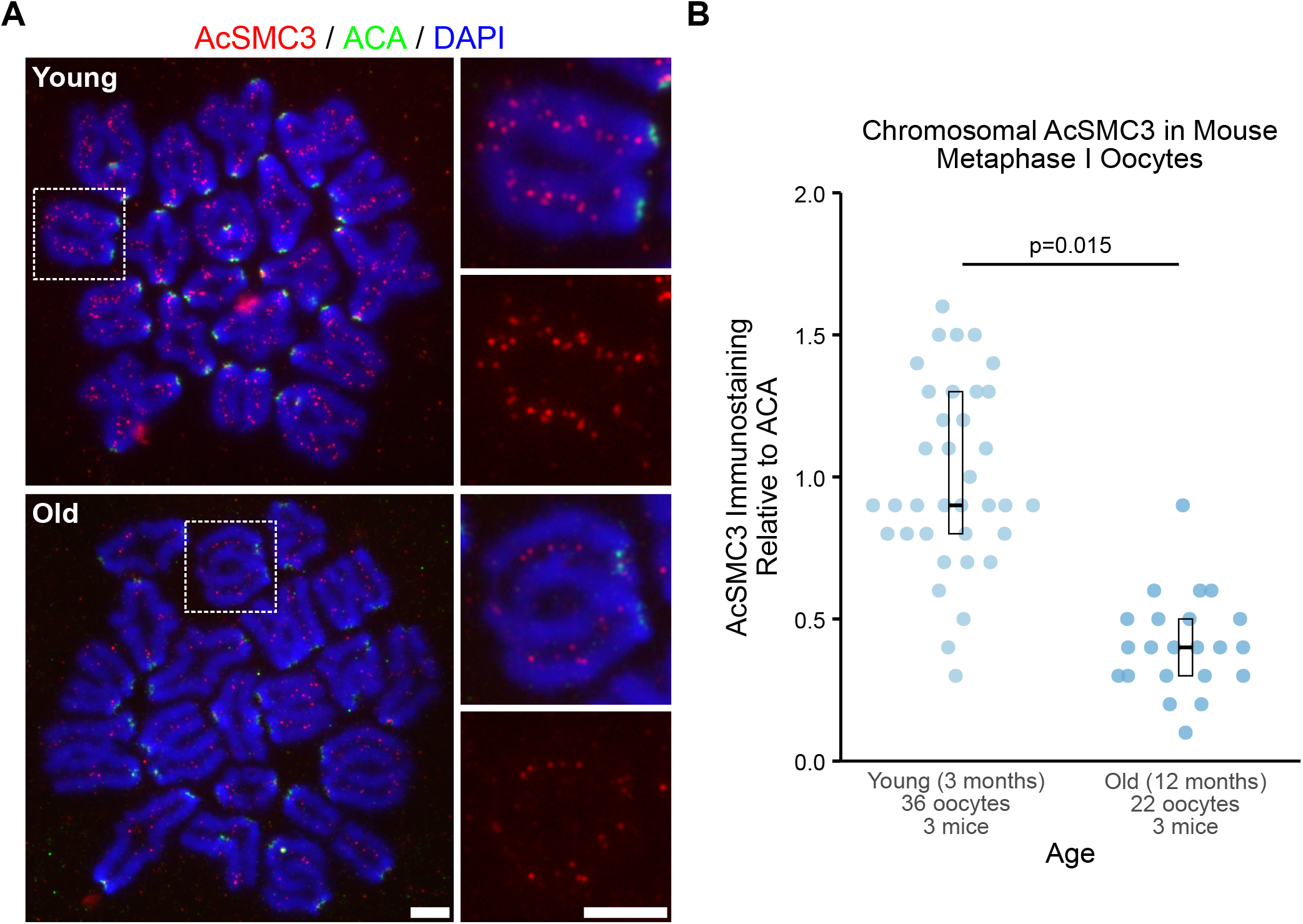
AcSMC3 is depleted from metaphase I mouse chromosomes from older mice. **A.** Immunostaining of chromosome spreads from metaphase I oocytes with antibodies against AcSMC3 (red) and anti-centromeric antibodies (ACA, green). DNA is counterstained with DAPI (blue). Dashed rectangles, higher magnification panels. Young and old mice were 3 and 12 months old respectively. Scale bars 5 µm. **B**. Quantification of AcSMC3 staining on chromosome spreads from metaphase I oocytes. AcSMC3 immunostaining intensity within each bivalent was measured relative to ACA, median relative AcSMC3 intensity for each oocyte is shown with a blue dot. Crossbars indicate the first, median and third quartiles; p values, linear mixed-effect models.

We next tested whether AcSMC3-containing cohesin was being lost while dictyate oocytes remain dormant within primordial follicles. Distinguishing non-chromosome-associated cohesin subunits, the cohesive subpopulation of chromosome-associated cohesin and other subpopulations of chromosome-associated cohesin involved in different aspects of chromosome biology^30^ is potentially difficult when immunostaining tissue sections. However, in somatic cell types, the cohesive subpopulation of cohesin is acetylated on the SMC3 subunit after it has been loaded onto the chromosomes and is rapidly deacetylated once it is removed from chromosomes^51^, therefore anti-AcSMC3 immunostaining is likely highly enriched for chromosome-associated cohesin functionally mediating sister chromatid cohesion. Interestingly, nuclear anti-AcSMC3 immunostaining in primordial oocytes is significantly 1.8-fold lower in ovary sections from 12-month-old mice than 3-month-old mice (**Figure 5**). The magnitude of this change is similar to the difference we detected between anti-AcSMC3 staining of metaphase I oocyte chromosome spreads from young and old mice (**Figure 4**) and is consistent with our observations from FISH on dictyate stage primordial oocytes showing that sister chromatid separation increases with age at this stage of oogenesis (**Figure 1**). Thus, mouse dictyate oocytes in dormant primordial follicles appears to lose the AcSMC3-marked cohesive subpopulation of cohesin from their chromosomes as they age, and the reduced levels of AcSMC3-cohesin in older primordial oocytes likely persists through subsequent stages of oocyte development until the oocytes reach metaphase I.

**Figure 5.**
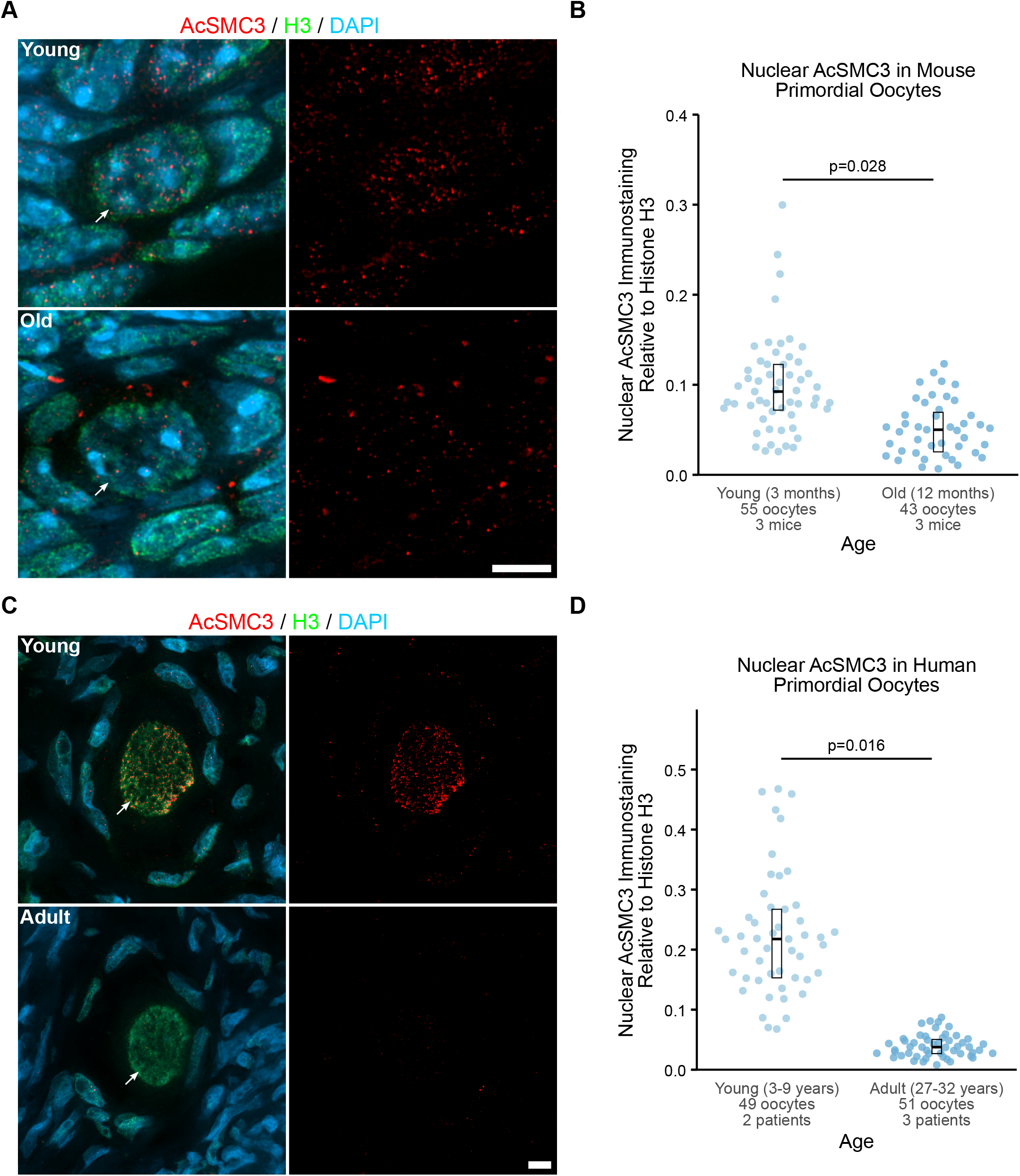
AcSMC3 is depleted from primordial oocytes with age in both mice and humans. **A, C**. Immunostaining of ovary sections from young (3-month-old) and old (12-month-old) mice (A), and young patients (3-9 years-old) and adult (27-32 years old) women (C). Anti-AcSMC3 staining is shown in red, anti-histone H3 in green, DNA is counterstained blue with DAPI. Images are 3D sum projections. Scale bars 5 µm. **B, D**. Quantification of nuclear AcSMC3 staining relative to histone H3 in young and old mouse (B) and young and adult human (D) primordial oocytes. Blue dot, oocyte relative AcSMC3 intensity; crossbars, first, median and third quartiles; p-values, linear mixed-effect models.

Similarly, anti-AcSMC3 staining of human ovary tissue sections showed that AcSMC3 is present in a thread-like pattern in the nucleus of human primordial oocytes, consistent with previous reports of cohesin immunostaining^24,39^. Strikingly, nuclear anti-AcSMC3 immunostaining in primordial oocytes from 27–32-year-old adult women is 5.8-fold lower than in 3-9-year-old girls (**Figure 5**). Thus, the cohesive AcSMC3-marked subpopulation of cohesin exhibits an age-dependent decline in human oocytes while they remain dormant at the primordial follicle stage. Furthermore, the age-dependent increase in chromatid separation that we detected by FISH in both mouse and human primordial oocytes appears to be associated with an age-dependent loss of the AcSMC3-marked cohesive subpopulation of cohesin in these cells in both these species. We propose that this common age-dependent loss of sister chromatid cohesion while oocytes remain dormant at the primordial follicle stage contributes to the age-dependent increase in meiotic chromosome mis-segregation and oocyte aneuploidy observed in both mice and humans.

## Discussion

### Chromosome ageing in mammalian oocytes

Our findings that age affects chromatid separation and cohesive AcSMC3-marked cohesin while mouse and human oocytes remain dormant in primordial follicles aligns well with previous mouse studies showing that chromosome-associated REC8-cohesin and chromatid cohesion have become depleted in older metaphase I oocytes^7,9,10^. However, age-dependent depletion of REC8-cohesin from human metaphase I oocyte chromosomes has not yet been detected^24^. Though increasing the age range and/or sample size may allow age-dependent depletion of REC8 to be detected in human metaphase I oocyte chromosomes, it is also possible that the REC8-cohesin responds differently to age in mice and humans, potentially reflecting species-specific differences in ovarian ageing^8^. It is possible that there are multiple mechanisms contributing to age-dependent depletion of chromosome-associated cohesins across multiple stages of mammalian oogenesis^8,12–14^, with more interspecies commonality in the effect of ageing on AcSMC3-cohesin than on REC8-cohesin. The age-dependent changes in chromatid separation and AcSMC3-marked cohesive cohesin described here potentially contribute directly to the high rates of premature sister chromatid separation and reverse segregation in oocytes from women of advanced maternal age^5^. Better understanding of how the AcSMC3-containing subpopulation of cohesin is maintained and regulated in postnatal oocytes, particularly those in the dormant primordial follicle stage, may better illuminate how ageing impacts on meiotic chromosome segregation in mammals.

Our data suggests that at least some age-dependent loss of chromatid cohesion occurs in fertile women during the child-bearing years and may directly contribute to the high frequency of inherited aneuploidy in human conceptions^1–6^. The duration of the primordial follicle stage is directly related to maternal age, therefore it seems plausible that age-dependent effects can occur during this stage of oogenesis. However, age-related changes in the ovarian somatic environment could also influence meiotic chromosomes during later stages of folliculogenesis^14,15^. Differences in chromatin organisation^52^ may complicate direct comparisons of chromatid separation between follicular stages using FISH, therefore we cannot exclude the possibility that additional chromatid cohesion is lost at later stages of folliculogenesis. In mouse oogenesis, depletion of AcSMC3 during the primordial follicle stage appears to account for most, though perhaps not all, of the loss of AcSMC3 observed in 1-year-old oocytes at metaphase I.

### Different chromosomal regions are differentially affected by ageing in mammalian oocytes

This study used six different FISH probes across two species to assess chromatid separation. While all four telomere-proximal probes detected age-dependent changes in primordial follicles, neither centromere-proximal probe showed a strong age-dependent increase in chromatid separation. This is consistent with previous data suggesting that centromeric cohesion is protected during meiosis I^33,48–50^. Increased protection of centromeric cohesion might be expected to result in more primordial oocytes containing 2 foci for centromere-proximal locations, however our FISH data do not fully support this. Furthermore, total inter-foci distances in 2-foci primordial oocytes, which likely represent inter-homolog distances, were significantly less for centromere-proximal probes than for telomere-proximal probes on the same chromosome. Further work is needed to understand whether the behaviour of these centromere-proximal regions does relate to their proximity to centromeres rather than other features at these chromosomal regions, including chiasma, that might influence chromatid separation. In addition, although all three human telomere-proximal probes exhibit age-dependent increases in chromatid separation in human primordial oocytes, there are some differences between them. These differences may represent different chromosomal regions being at different points in a continuum where ageing first affects separation of sister chromatids, then separation of homologs. Further experiments investigating which regions of primordial oocyte chromosomes are enriched for cohesive cohesin may allow a more directed approach to targeting FISH probes to chromosomal regions that might be more or less sensitive to age-dependent changes. Regional variation in cohesive cohesin, or sensitivity to ageing, may contribute to some of the differences in chromatid behaviour in this study, and possibly also to susceptibility of different chromosomes to some types of age-dependent meiotic chromosome mis-segregation events^5^.

### Evolving species differences in chronological time during oogenesis

The time frame over which ageing affects primordial oocytes differs significantly between mouse and human, with the reproductive life-span of mice lasting from approximately 6 weeks to 15 months after birth, and that of humans from approximately 15 to 50 years^53^. If the cohesion exhaustion model for oocyte ageing^12^ applies to both species, then the amount of cohesive cohesin initially established on meiotic chromosomes and/or the rate of its loss in primordial oocytes must be different. It is possible that more AcSMC3 is established on human oocyte chromosomes than mouse, though studies on fetal stages would be needed to investigate this further. However, assuming a constant depletion rate of AcSMC3 across the ages analysed in this study, the rate at which AcSMC3 immunostaining is depleted in human primordial oocytes is approximately 10-fold slower than in mouse. Even if the rate of AcSMC3 depletion in human primordial oocytes changes between pre-pubertal and post-pubertal stages, the rate in human primordial oocytes is still at least approximately 5-fold slower than in mouse. We propose that the rate of AcSMC3 depletion in primordial oocytes has been evolutionarily tuned to align cohesive cohesin levels chronologically with reproductive lifespan.

Interspecies differences in developmental tempo between mice and humans in neural progenitor cells and in pre-somitic mesoderm are determined largely by differences in protein stability^54,55^. In these contexts, proteins are more stable in a human cellular environment, with proteins in human neural progenitor cells generally having a 2-2.5-fold longer half-life than in mouse^54,55^. It is possible that the age-dependent depletion of cohesive cohesin in primordial oocytes partly reflects proteostatic degradation of chromosome-associated cohesins which cannot be replaced at this stage of oogenesis^26,35,36^, with general interspecies differences in protein stability contributing to the developmental allochrony. Identification and characterisation of the molecular pathways that impact on the stability of cohesive cohesin in primordial oocytes may provide insights into how evolution has aligned the chronology of cohesin maintenance with that of oogenesis and reproduction in different species, and identification of potential therapeutic targets to delay or prevent meiotic chromosome ageing in oocytes.

## Acknowledgements

We thank University of Edinburgh BioResearch and Veterinary Services and Institute of Genetics and Cancer Advanced Imaging Resource for support with the animal and imaging respectively, and Javier Caceres (University of Edinburgh) for comments on the manuscript. This study was supported by Medical Research Council University Unit awards (MC_UU_00007/6, MC_UU_00035/3, MC_UU_00035/7) (MB, AM, HB, SB, IRA), Medical Research Council grant (G1100357) (RAA), a Wellcome Trust Collaborator Award (215625) (EET and RAA), an MRC Doctoral Training Programme studentship (MB), and a Royal Thai Government studentship (AM).

## Author contributions

AM developed the FISH approach for mouse ovary tissue and acquired the mouse FISH data, MB applied the FISH approach to human ovary tissue and acquired and analysed the human FISH data, and the human and the mouse AcSMC3 immunostaining data. SB provided FISH expertise. HB performed data analysis and provided statistical advice. VS, YLO, RAA and EET provided human ovary tissue and human follicle staging expertise. IRA analysed the mouse FISH data. IRA and EET conceptualised, developed and managed the study.

## Materials and Methods

### Mouse Ovary Tissue

Animal use was in accordance with UK Home Office legislation and approved through project licence PPL P39007F29. Adult C57BL/6J mice (Charles River) were maintained until the required age according to institutional guidelines (University of Edinburgh, Bioresearch and Veterinary Services) and culled by cervical dislocation. Ovaries were dissected into PBS then fixed in 4% paraformaldehyde in PBS at 4 °C overnight. Ovaries were washed in PBS, then dehydrated through a 70% to 100% ethanol series, cleared in xylene, and infiltrated with paraffin wax at 58 °C in a Tissue-Tek VIP vacuum infiltration VIP tissue processor (Sakura). Tissue blocks were then sectioned at 8 µm thickness using a microtome, collected on Superfrost Plus slides, and dried at 50 °C overnight.

### Human Ovary Tissue

Ovary tissue was obtained from patients aged 3, 5, 9 and 14 years who were undergoing removal of ovarian cortex for fertility preservation prior to chemotherapy or radiotherapy for malignant disease. Samples of adult ovarian tissue from healthy women aged 25-40 years were obtained with informed consent from women undergoing elective Caesarean section. Protocols for both fertility preservation and donation for research had Ethical Committee approval (LREC 06/S1103/26 and 10/S1101/216/SS/0114), and all patients and/or their parents gave informed consent in writing.

Ovarian biopsies were collected in theatre and placed in pre-warmed dissection medium (Leibovitz L-15 supplemented with 2 mM glutamine (Life Technologies), 2 mM sodium pyruvate, 3 mg/mL human serum albumin, 75 µg/mL penicillin G and 50 µg/mL streptomycin (all from Sigma-Aldrich)). The ovarian cortex was dissected from the medulla and divided into fragments as previously described ^56^ using a scalpel blade and fine forceps and placed in fresh warmed dissection medium. Cortical strips were fixed in neutral-buffered formalin. After fixation for 24 h, individual fragments were dehydrated through graded alcohols and cleared in cedar wood oil before being embedded in paraffin wax and serially sectioned at 6 µm thickness.

### Fluorescence in situ hybridisation probe preparation

Fosmid clones (**Table 1**) were obtained from BACPAC Genomics and isolated from bacterial cultures in L-broth containing chloramphenicol using alkaline lysis. Fosmid DNA was directly labelled by nick translation using ChromaTide Alexa Fluor 594-5-dUTP (Life Technologies) or Green 496 dUTP (Enzo Life Sciences), DNaseI (Invitrogen) and DNA polymerase I (Life Technologies) ^57^. Labelled probes were purified through Quick Spin (Roche) or Qiaquick PCR Purification columns (Qiagen). Chromosome paints directly labelled with fluorophore were purchased from MetaSystems Probes.

**Table 1.**
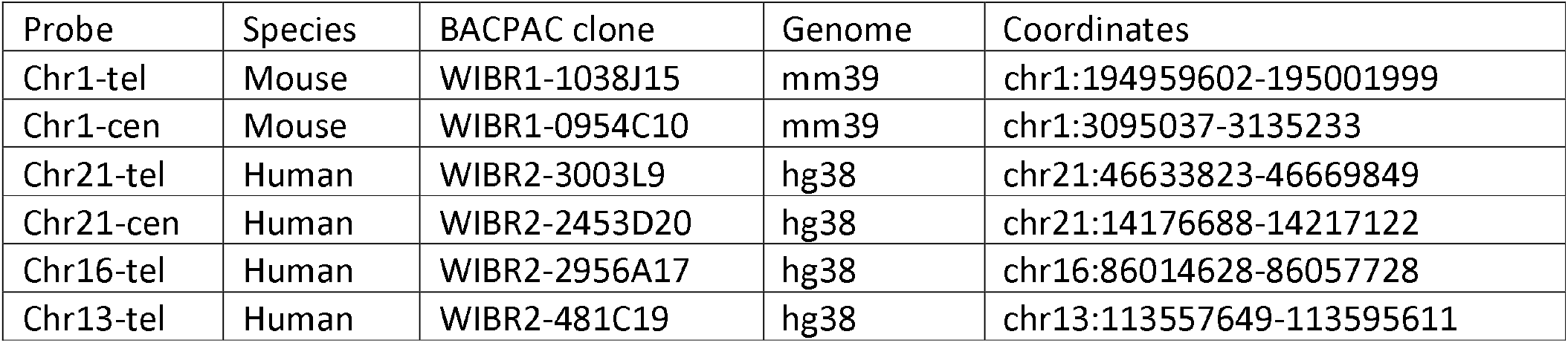
FISH probes used in this study.

### Fluorescence in situ hybridisation on ovary tissue sections and chromosome spreads

Ovary tissue slides were dewaxed with three washes in xylene for 5 minutes each, rehydrated through a 100% to 30% ethanol series then washed in dH_2_O. Antigen retrieval was performed by boiling in 10 mM sodium citrate pH 6 in a microwave for 45 minutes. Slides were subsequently allowed to cool at room temperature, warmed in 2×SSC at 75 °C for 5 minutes, denatured in 70% formamide, 2×SSC, pH 7.5 at 75 °C for 3 minutes, then snap-cooled in ice-cold 70% ethanol for 3 minutes. Slides were then dehydrated through 90% and 100% ethanol for 2-3 minutes each, and air-dried.

For each ovary tissue section slide, 130-160 ng each labelled fosmid probe, 8 µg mouse or human Cot-1 DNA as appropriate (Invitrogen) and 5-10 µg sonicated salmon sperm DNA (Invitrogen) was ethanol precipitated then resuspended in 40 µl hybridisation buffer (50% deionised formamide, 10% dextran sulphate and 1% Tween-20 in 2×SSC). The hybridisation mix was denatured at 70-75 °C for 5 minutes, pre-annealed for at least 15 minutes at 37 °C, then hybridised under a sealed coverslip at 37 °C from overnight to up to 2 days. The coverslips were then removed and the slides washed through four washes with 2×SSC at 45 °C for 3 minutes each, four washes with 0.1×SSC at 60 °C for 3 minutes each, and one wash with 4×SSC with 0.1% Tween-20 for 5 minutes ^57^. In some experiments 2 µg/mL DAPI was added to this last wash. Finally, the slides were mounted with Vectashield containing DAPI (Vector Laboratories) and sealed with clear nail polish.

FISH on ovary tissue sections was imaged in 3D by acquiring z-stacked images using a Coolsnap HQ II camera (Photometrics) or Prime BSI sCMOS camera (Teledyne Photometrics) mounted on a AxioImager A1 or Z2 fluorescent microscope (Carl Zeiss) equipped with a PIFOC P-721 objective scanning system (Physik Instrumente). Mouse ovary images were deconvolved using the Richardson-Lucy method using NIS offline deconvolution software (Nikon), human ovary images were deconvolved using iterative deconvolution with calculated PSF in Zen Acquisition software (Carl Zeiss). Analysis of FISH images and calculation of 3D distances between FISH signals was performed using the ImageJ 3D suite ^58^ to find the x, y and z co-ordinates of the centre of mass for each FISH focus. 3D distances between foci centres of mass were then calculated using Pythagoras theorem. 3D FISH signals were often located in different optical z sections. To represent the mouse data in 2D, maximum z projections of optical sections containing the deconvolved FISH signal were generated in FIJI ^59^, and superimposed on a single z section that contained the oocyte nucleus and allowed classification of the follicle (primordial follicles have flattened squamous granulosa cells surrounding the oocyte, primary follicles have a single layer of cuboidal epithelial granulosa cells surrounding the oocyte). For human ovary FISH, images represent maximum z projections of deconvolved FISH and DAPI signals, primordial follicles were identified as having flattened squamous granulosa cells surrounding the oocyte, primary follicles were identified as having a single layer of cuboidal epithelial granulosa cells surrounding the oocyte. Transitory follicles that had a mixture of squamous and epithelial granulosa cells were not included in these analyses.

Statistical analyses of the proportion of oocytes containing two, three or four discrete FISH foci was performed by generating baseline-category logit models for foci number using individual patients or mice as a random nuisance factor, and comparing Akaike’s Information Criterion (AIC) to determine the relative likelihood between a null model not incorporating age and a model incorporating age as an effect fitting the data. If models did not converge due to foci number categories segregating completely with an age group (**Figure 1D**), models were simplified by comparing oocytes containing two FISH foci to oocytes containing more than two FISH foci. For statistical analysis of distances or immunofluorescence staining intensities, linear mixed-effect models were built using the patient or mouse as a random nuisance factor. Pairwise distances between foci were separated into two populations using mclust^60^.

### Fluorescence in situ hybridisation on mitotic chromosome spreads

To obtain chromosome spreads from cultured cells, mouse embryonic stem cells (E14Tg2a) or human HEK293T cells were grown on gelatinised tissue culture plastic as described ^61^, trypsinised using 1% trypsin/0.4% EDTA (Sigma-Aldrich) and pelleted at 400*g* for 5 minutes. Cells were washed with PBS, pelleted, then resuspended in 10 mL hypotonic solution (mouse embryonic stem cells, 0.5% tri-sodium citrate, 0.25% potassium chloride; HEK293T cells, 0.5% tri-sodium citrate, 0.56% potassium chloride) and incubated for 10 minutes at room temperature. After pelleting the cells at 400*g* for 5 minutes, the cells were resuspended and fixed in 3:1 methanol-acetic acid for 5-10 minutes at room temperature. Cells were subsequently pelleted and washed with fixative at least three more times, dropped on clean microscope slides, and allowed to air-dry.

Hybridisation mix for FISH on mitotic chromosome spreads was prepared similarly to that described for ovary tissue sections except that half the amount of DNA was prepared in 15 µl of hybridisation buffer for each slide. Mitotic chromosome spread slides were incubated in 2×SSC with 100 µg/ml RNase A (Thermo Fischer Scientific) at 37 °C for 1 hour, then dehydrated through a 70% to 100% ethanol series. Slides were air-dried, then warmed at 70 °C in an oven for 5 minutes, then denatured, hybridised, washed and mounted as described for ovary tissue sections. FISH on mitotic chromosome spreads was imaged using the Coolsnap HQ II camera (Photometrics) mounted on a Zeiss Axioplan2 fluorescence microscope (Carl Zeiss). MicroManager software was used to capture images.

### Acetylated SMC3 immunostaining on mouse oocyte chromosome spreads

Female C57BL/6J mice were superovulated by hormonal injection of 0.1 mL 5IU pregnant mare serum gonadotrophin, then culled by cervical dislocation and ovaries collected in M2 media (Sigma) 42 hours later. Oocytes were released by piercing the ovaries repeatedly with a 25G needle, and germinal vesicle (GV)-stage oocytes selected based on oocyte shape, size and GV position. GV-stage oocytes were then cultured in pre-warmed M16 media (Sigma) at 37 °C, 5% CO2 for 2 hours and oocytes that had undergone GV breakdown selected and cultured for a further 5 hours to reach metaphase I. Preparation of metaphase I oocyte chromosome spreads and anti-AcSMC3 (Sigma-Aldrich, MABE1925, 1:500 dilution) and ACA (Antibodies Inc, 15-235-0001, 1:50 dilution) immunostaining performed as described ^40,62^. 2D epifluorescence images were acquired using an Orca Flash 4.0 cMOS camera (Hamamatsu) mounted on a AxioImager M2 fluorescence microscope with Plan-neofluor/apochromat objective lenses (Carl Zeiss) using Micromanager (Version 1.4) with scripts written by the IGC Imaging Facility. For immunofluorescence quantification, the bivalents in each image were manually traced in FIJI ^59^ using the DAPI channel, and the mean anti-AcSMC3 or ACA antibody signal above background per unit area determined for each bivalent. For each oocyte, the median normalised signal intensity across the twenty bivalents was then used to calculate the AcSMC3:ACA ratio. Image files were blinded by computer script prior to quantification.

### Acetylated SMC3 immunostaining on ovary sections

6 µm thick mouse or human ovary sections were dewaxed, rehydrated and antigen retrieval performed as described for in situ hybridisation. The slides were then blocked with 0.15% bovine serum albumin, 0.1% Tween-20 and 5% goat serum in PBS for 1 hour at room temperature, then incubated with primary antibodies (anti-AcSMC3, Sigma-Aldrich, MABE1073, 1:500 dilution; anti-Histone H3, Abcam, ab1791, 1:1000 dilution) for 3 hours at room temperature. After three washes for 5 minutes each in PBS, the slides were incubated with 20 µg/ml DAPI and Alexa Fluor-conjugated secondary antibodies (Invitrogen) for 1 hour at room temperature. The slides were then washed through three washes for 5 minutes each in PBS, treated with the TrueView Autofluorescence Quenching Kit (Vector Laboratories) for 2 minutes, then mounted with Vectashield containing DAPI (Vector Laboratories). 3D epifluorescence images were acquired using a Prime BSI sCMOS camera (Teledyne Photometrics Ltd) and a AxioImager Z2 fluorescence microscope (Carl Zeiss). Fluorescence illumination was provided by a Xylis LED light source (Excelitas Technologies) using single band filter cubes (Carl Zeiss, IDEX Health & Science). Images were deconvolved using an iterative deconvolution method with calculated PSF in Zeiss Zen 3.5 Acquisition software (Carl Zeiss UK, Cambridge, UK). The z-stacks were flattened using sum projections in FIJI ^59^, the outline of the nucleus manually traced using the DAPI channel, and the sum antibody signal intensity for AcSMC3 above background measured within the nucleus trace. This was normalised against the sum antibody signal intensity for Histone H3 above background within the nucleus trace. Image files were blinded by computer script prior to quantification.

## Supplemental information

Document S1. Supplementary Figures S1–S5.

Supplementary Figure S1. Validation of in situ hybridisation fosmid probes on mouse and human mitotic chromosome spreads

Supplementary Figure S2. Pairwise distances between chromatids increase with age at telomere-proximal regions in mouse and human primordial oocytes

Supplementary Figure S3. Fluorescence in situ hybridisation detects age-dependent changes in chromatid separation in mouse oocytes in primary follicles

Supplementary Figure S4. Age-dependent separation of the telomere-proximal region of chromosome 1 occurs during adulthood in human primordial and primary oocytes

Supplementary Figure S5. Ageing affects telomere-proximal chromatid separation specifically in oocytes but not granulosa cells in human primordial follicles

