## Supplementary Figures S1-S5 for "Chromosome Ageing Occurs at the Primordial Follicle Stage in both Mouse and Human Oocytes"

#### Document S1. Supplementary Figures S1–S5

#### Supplementary Figure S1. Validation of in situ hybridisation fosmid probes on mouse and human mitotic chromosome spreads

**A-B**. Left, ideogram showing the location of the mouse chromosome 1 telomere-proximal (A, Chr1-tel) and centromere-proximal (B, Chr1-cen) fosmid FISH probes (orange bars). Right, FISH on mouse embryonic stem cell metaphase chromosome spreads using a chromosome 1 paint (red), the Chr1-tel (**A**; green), and the Chr1-cen fosmid (**B**, green). DNA is stained with DAPI (blue); asterisks, centromeres (DAPI-dense regions). Scale bar 5 µm. **C**. FISH using both Chr1-tel (green) and Chr1-cen fosmid probes (red) on mouse embryonic stem cell metaphase chromosome spreads. DNA is stained with DAPI (blue); asterisks, centromeres. Scale bar 5 µm. **D-F.** Left, ideogram showing the location of the human fosmid probes (orange bars) used in this study. Right, FISH on human HEK293T metaphase chromosome spreads using the indicated chromosome paint (green) and fosmid probe (red). **D**, chromosome 13 telomere-proximal (Chr13-tel); **E**, chromosome 16 telomere-proximal (Chr16-tel); **F**, chromosome 21 telomere-proximal (Chr21-tel); **G**, chromosome 21 centromere-proximal (Chr21-cen). DNA is stained with DAPI (blue), asterisks, centromeres (constrictions between chromosome arms). Scale bar 5 µm.

#### Supplementary Figure S2. Pairwise distances between chromatids increase with age at telomere-proximal regions in mouse and human primordial oocytes

**A, B**. Density estimates of populations of pairwise distances between FISH foci for Chr1-tel in mouse primordial oocytes (A) and for Chr21-tel in human primordial oocytes (B). Data for pairwise distances were the same as that shown in Figure 1D (mouse) and Figure 2B (human). Mouse data were separated into younger (blue, 3-month and 6-month old) and older (red, 9-month old) age groups; young (blue) and old (red) age groups for human were as in Figure 2B. Pairwise distances for each age group were separated into two populations and density estimates for each population plotted (continuous and dashed lines). The mean pairwise distance for each population is indicated above its peak.

#### Supplementary Figure S3. Fluorescence in situ hybridisation detects age-dependent changes in chromatid separation in mouse oocytes in primary follicles

**A, C**. FISH for chromosome 1 telomere-proximal (Chr1-tel, A) and centromere-proximal (Chr1-cen, C) fosmids (red) in primary follicles in mouse ovary sections. FISH signals are 3D maximum projections. DNA is stained with DAPI (blue). Arrow, oocyte; arrowhead, cuboidal granulosa cell; dashed rectangle, insets (FISH channel only). Scale bar 5 µm. **B, D**. Quantification of Chr1-tel (B) and Chr1-cen (D) FISH foci in mouse primary oocytes. Left panel, distribution of oocytes with the indicated number of foci. RL, relative likelihood that a logit model not incorporating age fits the foci number distribution better than one that does. Right panel, total Chr1-tel (B) and Chr1-cen (D) inter-foci distance within 2-foci, 3-foci and 4-foci mouse primordial oocytes groups against age. Black lines, locally weighted scatterplot smoothing trends; p-values, linear mixed-effect models.

#### Supplementary Figure S4. Age-dependent separation of the telomere-proximal region of chromosome 1 occurs during adulthood in human primordial and primary oocytes

**A, B**. Quantification of the number of discrete chromosome 21 telomere-proximal FISH foci detected in adult human primordial oocytes (A) and adult human primary oocytes (B). Data shown are a subset (only adult women >18 years old) of the data shown in Figure 2B. Younger women, 25-32 years old; older women, 33-40 years old. RL, relative likelihood that a logit model not incorporating age fits the foci number distribution better than one that does. **C**. FISH for Chr21 telomere-proximal (Chr21-tel, A) and Chr21 centromere-proximal (Chr21-cen, C) fosmid probes (red) in primary follicles within human ovary sections. DNA is counterstained with DAPI (blue). Images are 3D maximum projections. Arrow, oocyte; arrowhead, cuboidal granulosa cell; asterisks, non-specific signal; dashed rectangle, insets (FISH channel only). Scale bar 5 µm.

#### Supplementary Figure S5. Ageing affects telomere-proximal chromatid separation specifically in oocytes but not granulosa cells in human primordial follicles

**A**. Quantification of chromosome 21 telomere-proximal (Chr21-tel) inter-foci distances in granulosa cells and in 3-foci oocytes in human primordial follicles. FISH foci coordinates for the oocyte data are the same as used to generate the total inter-foci distance per oocyte shown in Figure 2. Young patients, 3-14 years old; adult women, 25-36 years old. p-values, linear mixed-effect models. **B**. Quantification of chromosome 16 telomere-proximal (Chr16-tel) inter-foci distances in granulosa cells and in 2-foci, 3-foci and 4-foci oocytes in human primordial follicles. FISH foci coordinates for the oocyte data are the same as used to generate the total inter-foci distance per oocyte shown in Figure 3. Young patients, 3-14 years old; adult women, 25-36 years old. p-values, linear mixed-effect models.

**Bui et al., Supplementary Figure S1**

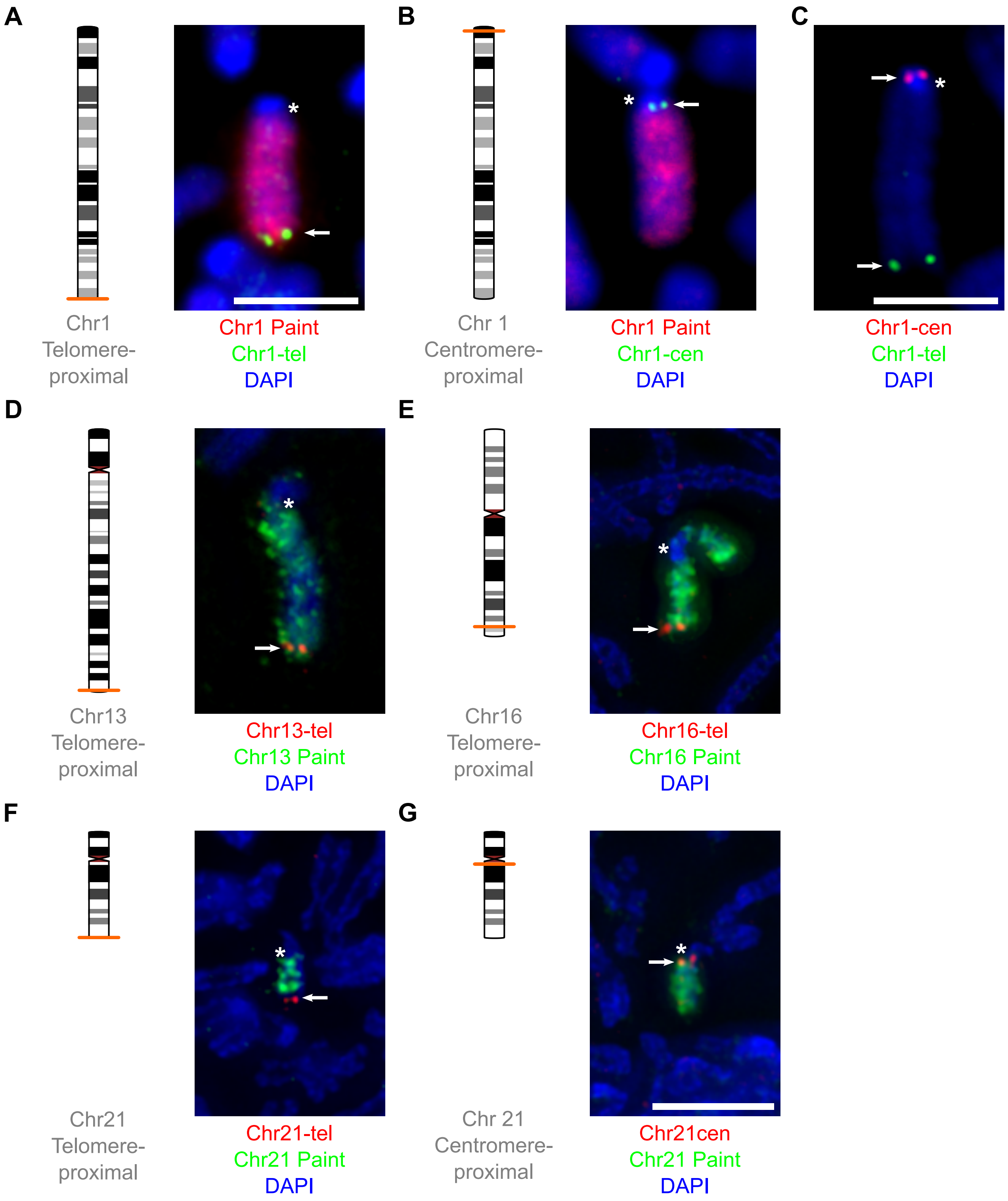

### Bui et al., Supplementary Figure S2

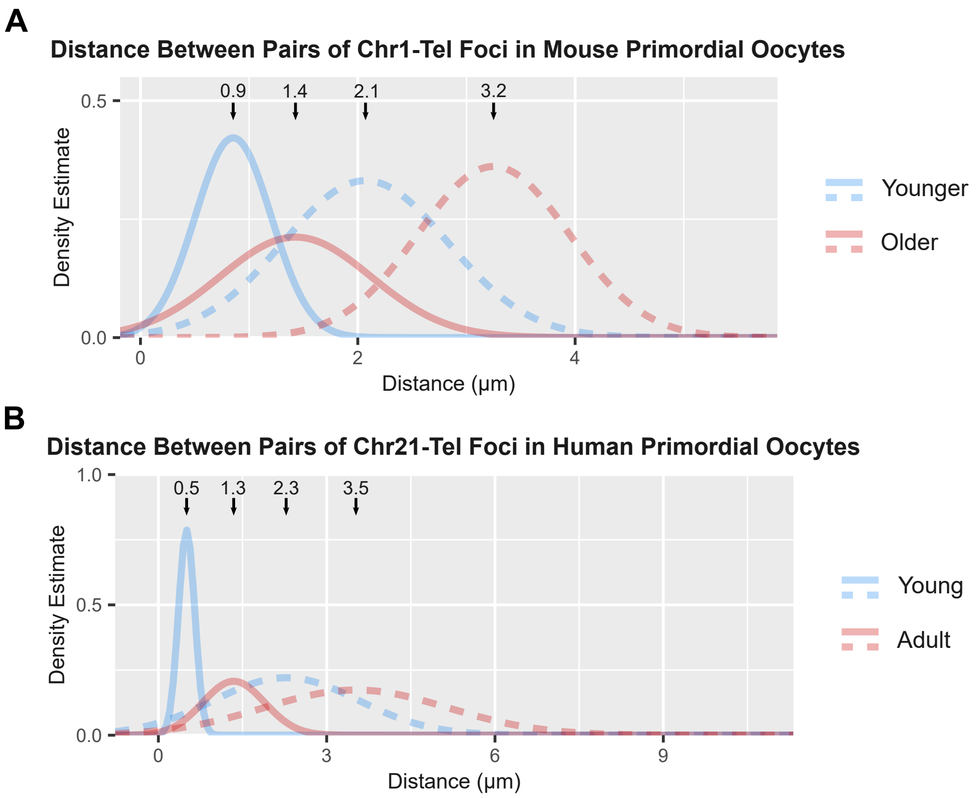

**Bui et al., Supplementary Figure S3**

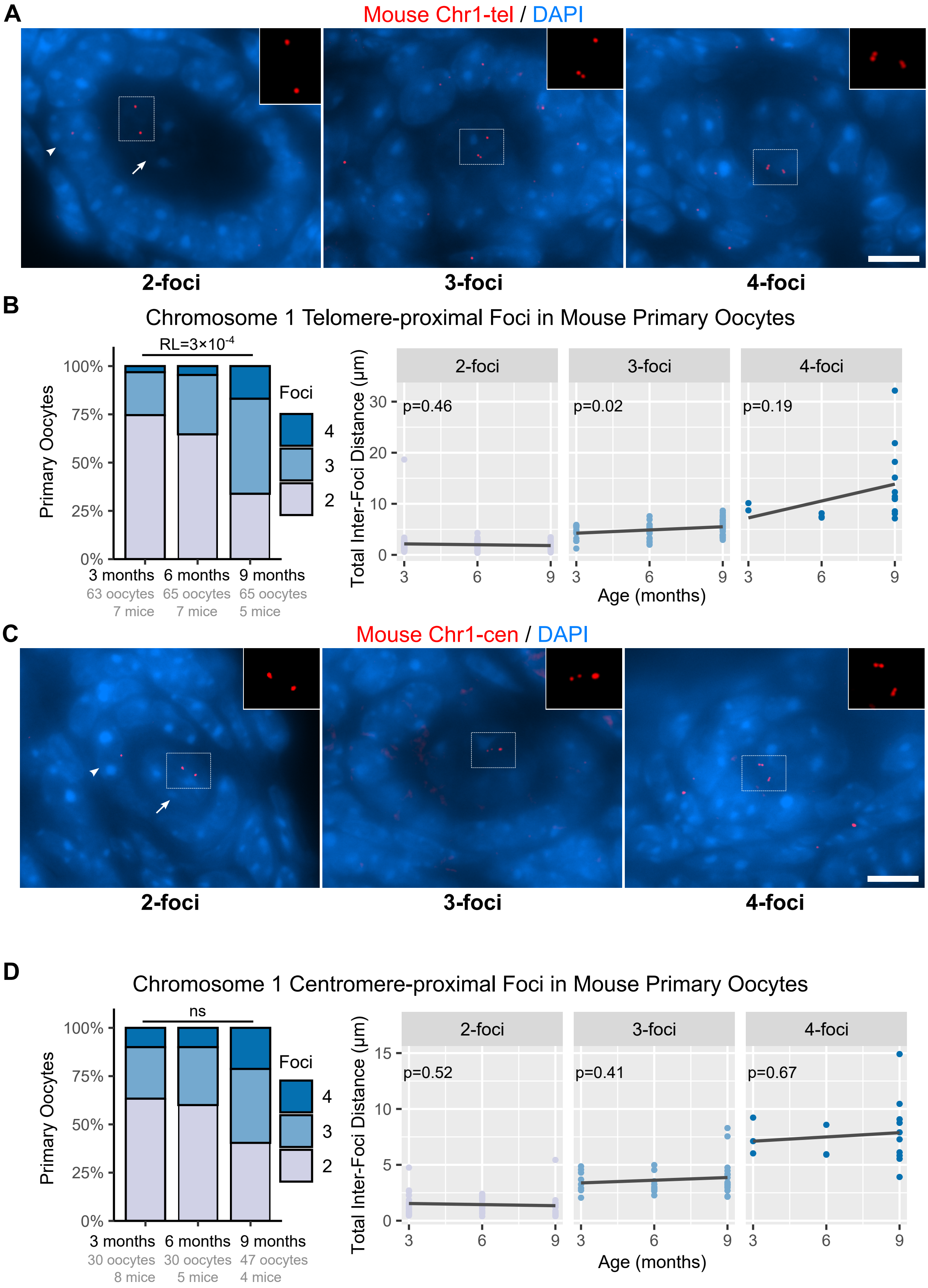

**Bui et al., Supplementary Figure S4**

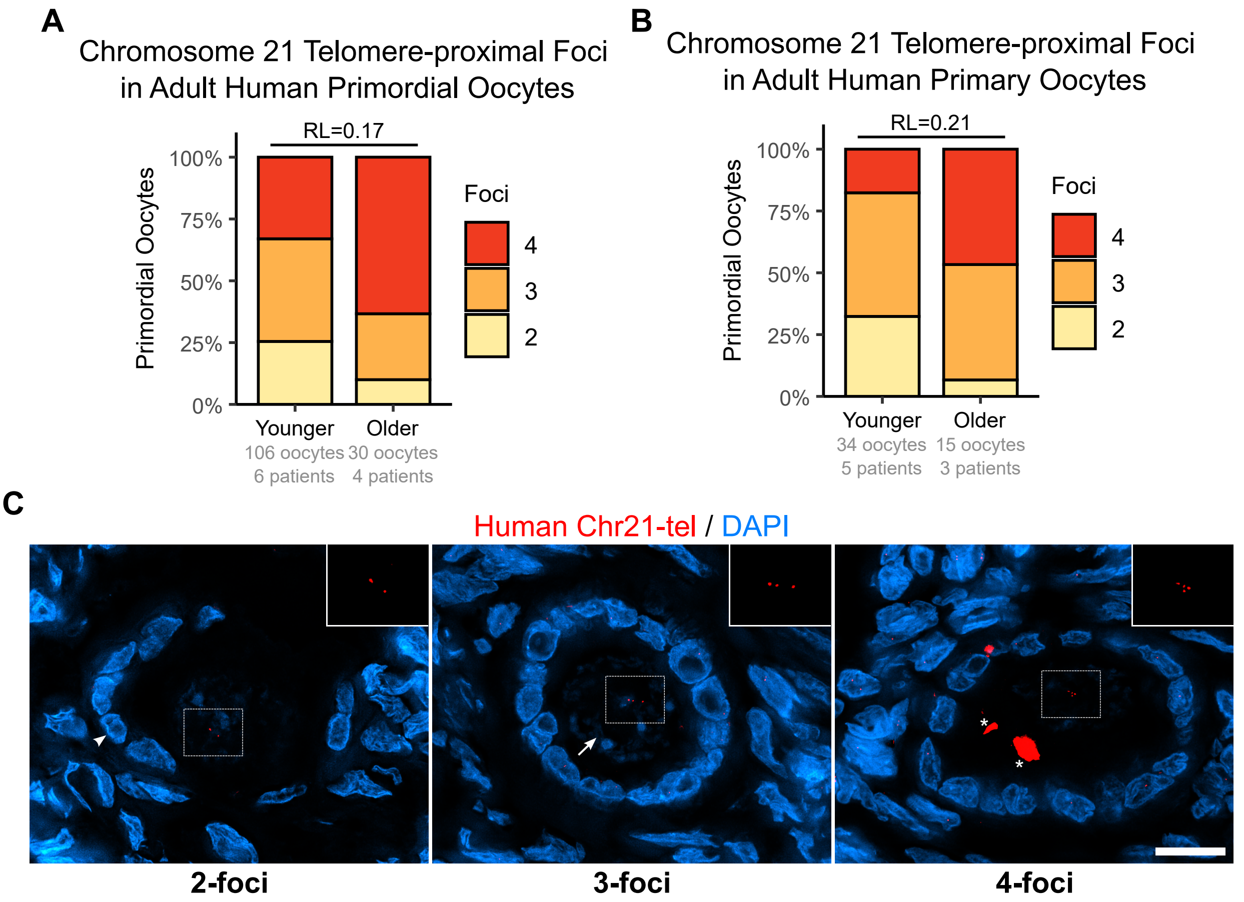

**A**

Chromosome 21 Telomere-proximal Foci in Human Primordial Follicles

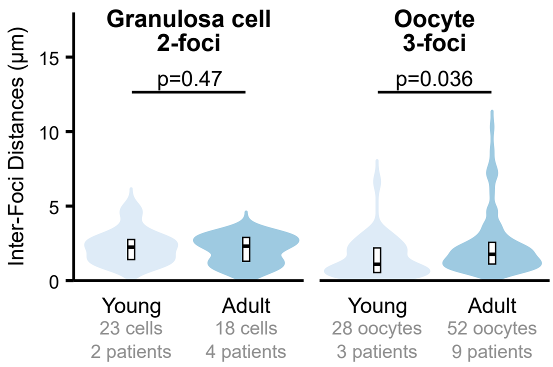


**B**

Chromosome 16 Telomere-proximal Foci in Human Primordial Follicles

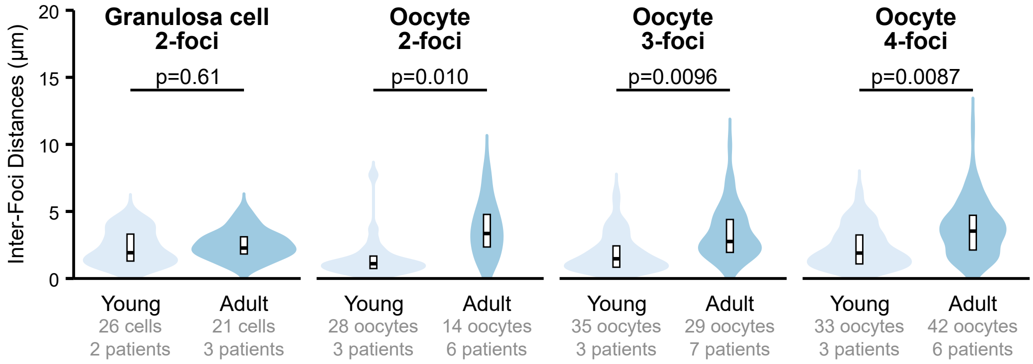
